# Addressing Inaccurate Nosology in Mental Health: A Multi Label Data Cleansing Approach for Detecting Label Noise from Structural Magnetic Resonance Imaging Data in Mood and Psychosis Disorders

**DOI:** 10.1101/2020.05.06.081521

**Authors:** Hooman Rokham, Godfrey Pearlson, Anees Abrol, Haleh Falakshahi, Sergey Plis, Vince D. Calhoun

## Abstract

**Background:** Mental health diagnostic approaches are seeking to identify biological markers to work alongside advanced machine learning approaches. It is difficult to identify a biological marker of disease when the traditional diagnostic labels themselves are not necessarily valid.

**Methods:** We worked with T1 structural magnetic resonance imaging data collected from individuals with mood and psychosis disorders from over 1400 individuals comprising healthy controls, psychosis patients and their unaffected first-degree relatives including 176 bipolar probands, 134 schizoaffective probands, 240 schizophrenia proband, 581 patients relatives and 362 controls. We assumed there might be noise in the diagnostic labeling process. We detected label noise by classifying the data multiple times using a support vector machine classifier, and then we flagged those individuals in which all classifiers unanimously mislabeled those subjects. Next, we assigned a new diagnostic label to these individuals, based on the biological data (MRI), using iterative data cleansing approach.

**Results:** Simulation results showed our method was highly accurate in identifying label noise. Both diagnostic and Biotype categories showed about 65% and 63% respectively of noisy labels with the largest amount of relabeling occurring between the healthy control and bipolar and schizophrenia disorder individuals as well as in the unaffected close relatives. The extraction of imaging features highlighted regional brain changes associated with each group.

**Conclusions:** This approach represents an initial step towards developing strategies that need not assume existing mental health diagnostic categories are always valid, but rather allows us to leverage this information while also acknowledging that there are misassignments.

## 1. INTRODUCTION

Psychiatry has struggled with identifying a biological basis for mental illness. Current categorization approaches, including the APA Diagnostic and Statistical Manual (DSM), are not entirely valid [1, 2, 3, 4]. Diagnosis of mental illness, such as schizophrenia and bipolar disorder, is typically based on unreliable symptom-based measures. It is also known that among different DSM diagnoses, there is a considerable overlap not only in their clinical symptoms but also in biological measures including disease risk genes, structural and functional brain measures, electrophysiology and cognitive functional deficits [5, 6, 7]. Additional challenges include the debate over the validity of additional “mixed” diagnostic categories such as schizoaffective disorder [5]. Unreliable self-report information regarding symptoms makes the process of diagnosis more complicated and leads to incorrect labeling. Understanding brain biomarkers in psychiatry may help to diagnose and treat mental disorders more effectively. Integrating clinical data with genomics and other patient information such as brain biomarkers help better define valid disease subtypes and/or categorize subjects more accurately and improve treatment outcomes [8]. The challenge is that applying biological classification while using traditional psychiatric diagnoses that lack inherent validity as ground truth is unlikely to prove productive.

Identifying and prioritizing diagnostic categorization is a major challenge in psychiatry which if not carried out correctly translates into inaccuracies in diagnostic labeling of biological data, such as medical imaging. Addressing these inaccuracies (which we refer to as label noise in a diagnostic classification problem setup) is an important topic of great interest that serves the ultimate goal of helping patients [9]. The application of artificial intelligence (AI) can be leveraged to help with this task and to achieve better results. Multiple studies have shown that there are significant differences between schizophrenia, bipolar and control regarding their structural imaging data especially in the cerebellum and temporal lobe regions [10] [11] [12] [13]. Among different forms of AI approaches, classification, the process of predicting the class of new samples has been broadly used in the realm of machine learning. To compute a prediction for an unseen sample, a supervised classifier algorithm first learns from data based on the provided labels. As such, the reliability of the dataset plays a vital role in the performance of the classification models. Therefore, if there is a label noise in the data set, the prediction accuracy will decrease. However, having a wholly pure and label noise-free dataset is unlikely in mental health, given the fact that current diagnostic strategies are inaccurate and diagnoses of questionable validity [14, 1]. Noise in this context refers to anything that obscures the relationship between the features of an instance and its class [1, 15]. In addition to the label noise, data quality issues e.g. low SNR, head motion, may corrupt the relationship between the features of an instance and its class as well. We performed quality control and preprocessing steps help to mitigate this, but we cannot rule it out completely. Previous work has shown how label noise may affect classification accuracy [1, 16]. Studies show that neural networks are robust in handling label noise in data [17]; however, deep learning models require large datasets [18] and the minimum amount of needed clean training data increases with an increase in the label noise level [17].

To improve categorization or nosology in psychiatry, it is essential to address the challenges in the existing categorization. One approach is to consider existing categories as ‘noisy’ (i.e., containing mislabeled samples) and develop approaches to eliminate or at least reduce the impact of this noise during a classification task. In this work, we identify cases where there is biological evidence that pushes against an existing categorization, that is, cases where gray matter data looks similar for subjects that are categorized differently or looks different for groups that fall within the same diagnostic category. Our proposed method includes using a voting approach to estimating individuals that are labeled (diagnosed) incorrectly, called class noise or label noise. Then, we relabeled noisy subjects with a new label such that subjects are more similar to each other in the new group. We then repeated these steps until we identified a specific acceptable amount of label noise in the dataset.

By applying our method, we identify shared features of existing categories in structural MRI data and regroup subjects that mislabeled or violate the assumption of homogeneity within groups and suggest the new labels similar to others which leads to having homogeneity within subjects of each category at the end. Results show that there are homogeneous subsets that fall both within and between existing categories and we suggest that future studies should focus more on these aspects of the available data.

## 2. MATERIALS AND METHODS

### 1. B-SNIP-1 DATASET

We analyzed the bipolar-schizophrenia network on intermediate phenotypes (B-SNIP-1) structural imaging (structural MRI) dataset [19, 20]. B-SNIP is multi-site NIH-funded consortium of investigators that collected multiple brain imaging and assessment measures for stable patients within three psychotic disorders (schizophrenia, schizo-affective disorder or bipolar disorder with psychosis). The B-SNIP-1 dataset used in this study included 912 subjects after quality control assessment. Structural MRI three-dimensional acquisitions were carried out on 3T scanners (GE Signa, Philips Achieva, Siemens Allegra, and Siemens Trio) [21] [22]. High resolution isotropic T1-weighted MP-RAGE sequences were acquired following the Alzheimer’s Disease Neuroimaging Initiative (ADNI) protocol [21, 23]. Subject demographics are reported in **Table 1**. More detail of B-SNIP dataset and the scanners described in supplemental.

**Table 1.**
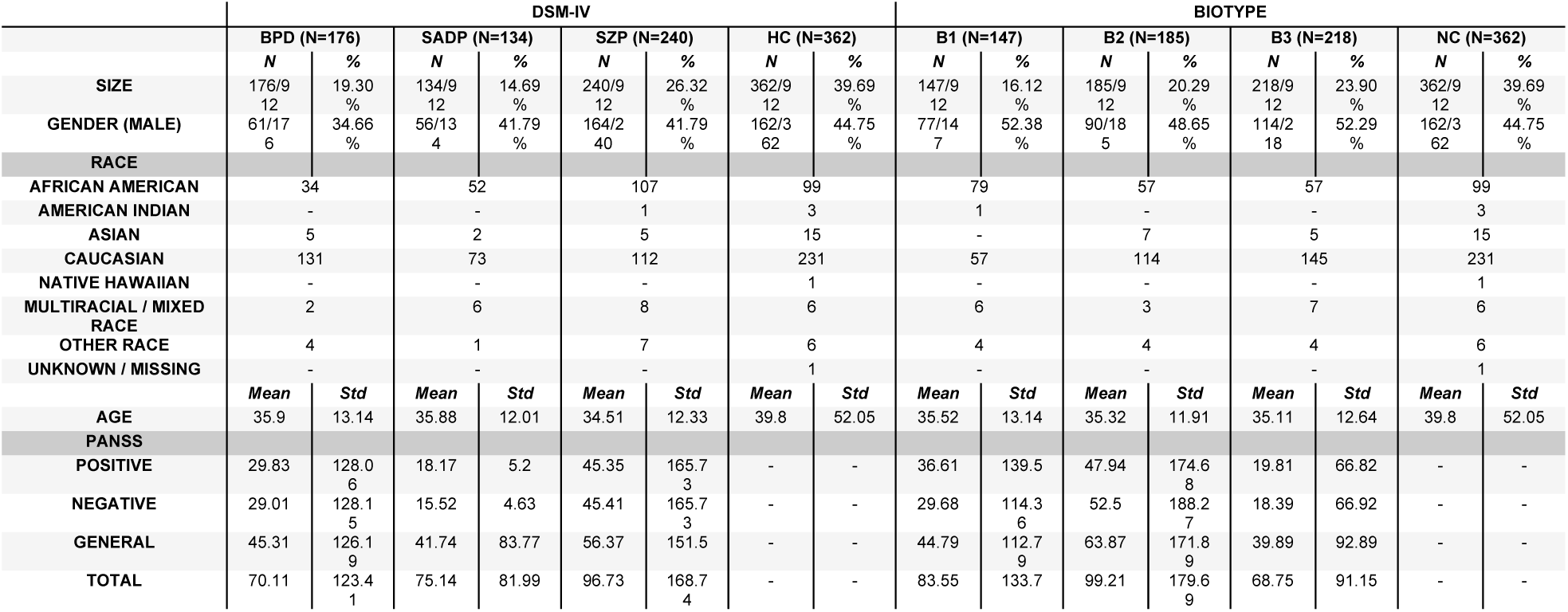
B-SNIP Dataset Demographic

### 2. PREPROCESSING

Structural MRI data (1mm isotropic MPRAGE) were collected from all individuals [19, 20]. Images were preprocessed in SPM (https://www.fil.ion.ucl.ac.uk/spm/) via a unified approach [24] that included tissue classification, bias correction, image registration, spatial normalization, and resliced to 2×2×2mm. The unsmoothed gray matter density (GMD) images were then correlated to the gray matter template to access segmentation outliers and outliers were corrected if possible or removed otherwise. We calculated the correlation of each gray matter segmentation map to the mean map and removed those that had lower correlation (<0.7). We have found these criteria to be very helpful in identifying problematic scans. We also centered and scaled the voxels of the subjects in each site to unit variance and covaried for each site separately.

### 3. METHOD DESCRIPTION

We used a novel classification-voting filtering method based on a data cleansing approach to eliminate label noise e.g. noisy, mislabeled subjects based on the structural MRI image dataset. This provided informative patterns for further investigation and for reassigning labels for identified mislabeled subjects based on the model. Our proposed model also provides suggested labels for noisy data sets. The method is based on computing *m* number of inner SVM classifiers that are trained and evaluated via cross-validation. We use super vector classifiers (SVC) from the scikit-learn python library [25] and we used a one-versus-all approach that has been implemented in the scikit-learn library in the way that it fits one classifier for each class against all the other classes. These *m* number of SVM models are then used to identify mislabeled subjects for different runs of cross-validation sets. Thus, each individual is classified *m*-times by SVM for *k* cross-validation loops totaling *m* × *k* classification votes. Based on the *m* × *k* predicted labels, we used consensus voting to determine if a given dataset includes a noisy label if all *m* × *k* number of voting labels are mislabeled. **Fig 1** presents a flow diagram of our method. In the following, we discuss more details regarding our proposed method.

**Fig 1.**
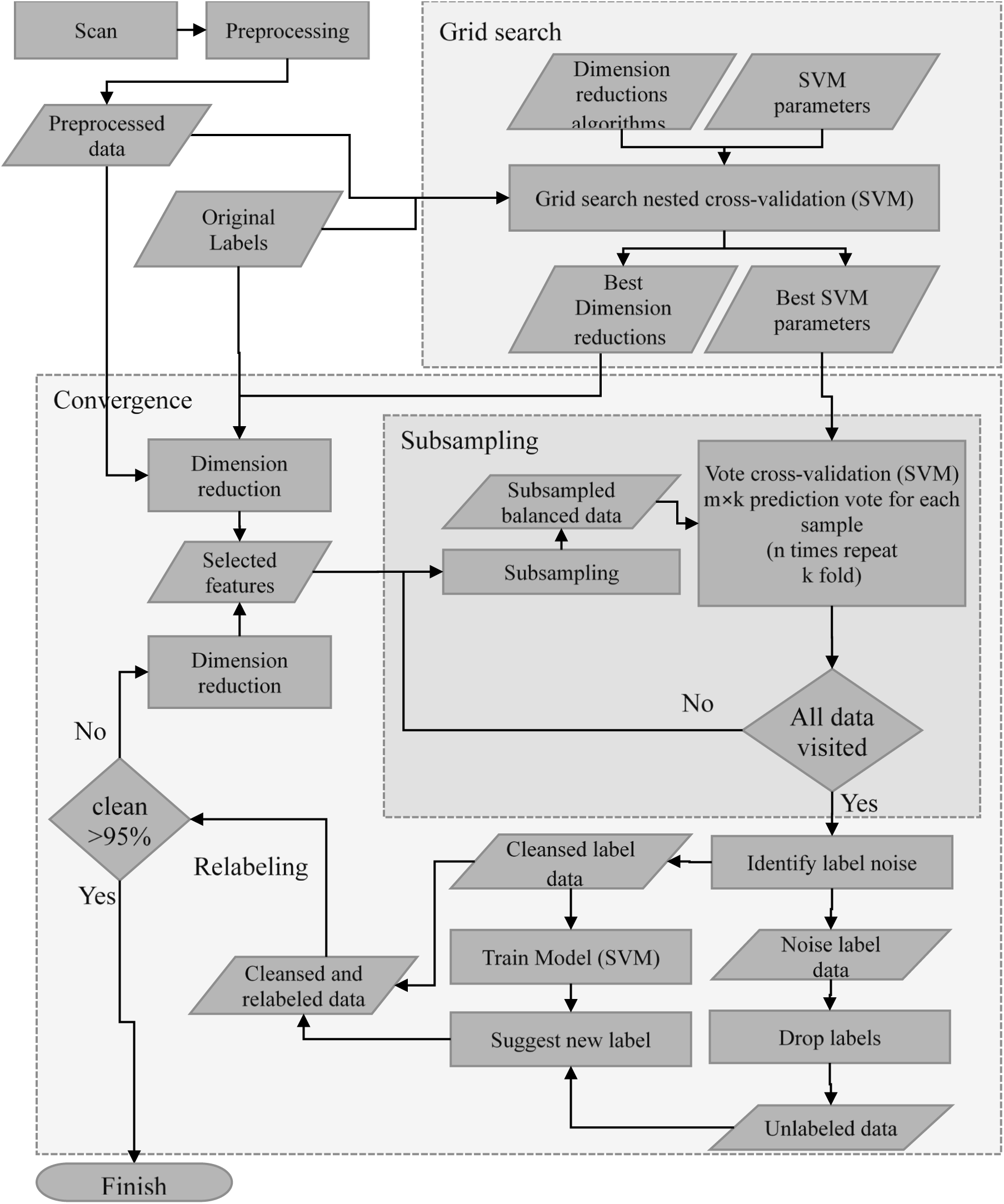
Visual summary and workflow of different aspects of the proposed method. Preprocessing was done using SPM on structural MRI data collected at multiple sites. Hyper-parameters optimization and parameter selection among choosing the best dimension reduction were done using grid search. Then univariate feature selection using ANOVA was selected as best approach and used for the dimension reduction approach, applied for dimension reduction on preprocessed data and subsampling was done on dataset to handle imbalanced classes. Cross validated classification/voting filtering was applied to resampled data. Then noisy labels were identified based on their votes, cleansed dataset (exclude noise subjects) and unlabeled dataset (noisy subjects with dropped labels) feed to supervised to evaluate and obtain the new suggested labels. Combining cleansed data and data with new suggested labels make cleansed and relabeled data. For generalization, using a convergence approach, subsampling, cross-validated classification voting filtering and relabeling performed iteratively on cleansed and new labeled data till we identified a specific acceptable amount of label noise in the dataset.

#### Simulation

Since there is no ground truth in the neuroimaging data, we first evaluate our method on handwritten digit images dataset, which enables us to compare our method on data for which there is well-defined ground truth. We introduce label noise to the dataset by shuffling a proportion of labels of instances randomly. We add different amounts of label noise and evaluate the model. Simulation studies described in more detail in supplemental.

#### Grid search

This approach seeks the hyper-parameter space through cross-validation and proposes the best candidate among possible hyper-parameters values. Grid search and the parameters’ space described in supplemental.

#### Dimension reduction and feature extraction

The curse-of-dimensionality comes with high dimensional neuroimaging scan data where the number of features (voxels) is significantly larger than the number of samples (brain images) [26, 27]. This may lead to overfitting and a lack of generalizability. We used univariate feature selection and computed a univariate ANOVA F-value between groups based on their assigned labels and used this to select the 100 best feature vectors for each cleansing iteration.

#### Imbalanced classes and subsampling

To address imbalanced classes we used a random under-sampling approach for handling imbalanced classes by resampling the majority class randomly and uniformly. This was done repeatedly until all the instances of the majority classes were visited at least one time.

#### Data cleansing classification/voting filtering

We obtain votes using cross-validated classification. Using the consensus voting approach, we considered individuals as noisy if all votes are inaccurate labels. Based on this, we keep all the data but remove the labels for noisy subjects and consider them to be an unlabeled sample for relabeling steps.

#### Relabeling using classification

We then trained the SVM model with cleansed data and predicted a new label for noisy subjects. The classification relabeling approach assumes that when two samples in a high-density region are similar then their output classification should be similar too. We also evaluated our relabeling dataset using another model, a deep learning residual model, to elucidate the effect of label noise on the accuracy of the predictive model by comparing accuracy before and after a data cleansing classification filtering method. The deep model described in more detail in supplementary and **Fig S3**.

#### Convergence

We repeated the process from subsampling step to relabeling step iteratively with new suggested labeled till the number of label noise identified in filtering step reach to an ad-hoc threshold (e.g. 5%). In each iteration, subsampling, classification voting filtering and relabeling perform on the cleansed and relabeled datasets (i.e. the entire dataset with original labels for cleansed data and newly assigned labels for label noise subjects). By using this iterative step, label noises gradually are detected and the performance of the model increases. Combining this iteration and randomly chosen partition for training and validating increase generalization of our approach.

#### Visualization

We performed voxelwise t-tests on gray matter maps to highlight brain regions where different groups differed significantly. For each pair of groups, we ran statistical t-tests on the original non-zero vectors of features (voxels) and then used the false discovery rate (FDR) correction for multiple comparisons. We identified those features that were significantly different between each pair of groups and created images only containing means of larger group of those features. We also plotted the dataset before and after cleansing on a 2D plot using 2D-tSTE [28].

## 3 RESULTS

### B-SNIP MRI Analysis

After validating our approach on simulation data (result of simulation in supplemental) we applied our method on the B-SNIP dataset. For the B-SNIP data, we used a univariate ANOVA F-value to select the 100 best features among 2,122,945 voxels for 912 individuals. Features extracted were most often found in amygdala, cerebellum and insula regions bilaterally. Some additional features which recurred included lingual, occipital, frontal, temporal, fusiform, para hippocampal and hippocampus regions. This was done for each type of labelling approach (Biotype and DSM). To handle imbalanced classes, we used the subsampling approach described previously.

Using a grid search approach, the SVM classifier with RBF kernel and penalty parameter 10 and kernel coefficient 0.04 was selected as the best model with optimized hyper-parameters for the B-SNIP dataset. The average accuracy of classification filtering is shown in **Fig 2**. The average SVM classifier cross validated accuracy increased from 0.38 to 0.89 for both Biotype and DSM-IV categories using consensus voting.

**Figure 2.**
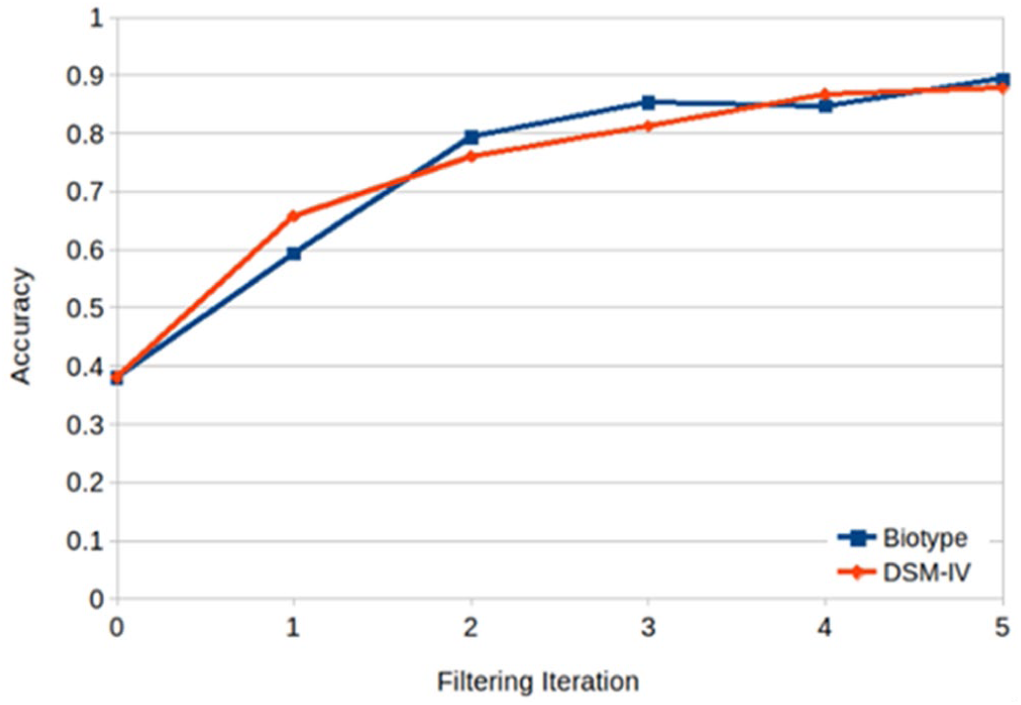
Average accuracy of classification filtering cross validation part after 5 iterations. In each iteration, label noise is identified, and new labels are assigned based on similarities between them and cleansed instances. The performance of the model increases after each iteration.

Among 912 subjects, 573 subjects (63%) were identified as label noise in the Biotype category and 601subjects (65%) were identified as label noise in the DSM-IV category. 452 out of 912 labels were identified as noisy for both Biotype and DSM-IV. Table 2 shows the different proportion and number of noisy labels found after 5 iterations of cross validation classification filtering and reveals convergence at approximately 90% of data has been cleansed. At each iteration we removed the labels for the noisy datasets (retaining the labels for the clean datasets) and fit an SVM model to predict new suggested labels. Fig 3 shows the heatmaps of those identified label noise in different types of labels. Also using the fit model, we predict the label for the relatives. 435 out of 581 relatives are labeled as Biotype B3 and normal control in the Biotype category and all of 581 are labeled as controls in the DSM-IV category.

**Table 2.** Shared label noise using consensus voting

| Consensus Biotype |  |  |  |  |  |  |  | Consensus DSM-IV |  |  |  |  |  |  |  |
| --- | --- | --- | --- | --- | --- | --- | --- | --- | --- | --- | --- | --- | --- | --- | --- |
| Clean:460 |  |  |  | Noise: 452 |  |  |  | Clean: 460 |  |  |  | Noise: 452 |  |  |  |
| B1 |  | B2 |  | B3 |  | NC |  | BPD |  | SADP |  | SZP |  | HC |  |
| Clean | Noise | Clean | Noise | Clean | Noise | Clean | Noise | Clean | Noise | Clean | Noise | Clean | Noise | Clean | Noise |
| 63 | 84 | 78 | 107 | 101 | 117 | 218 | 144 | 89 | 87 | 69 | 65 | 84 | 156 | 218 | 144 |

**Figure 3.**
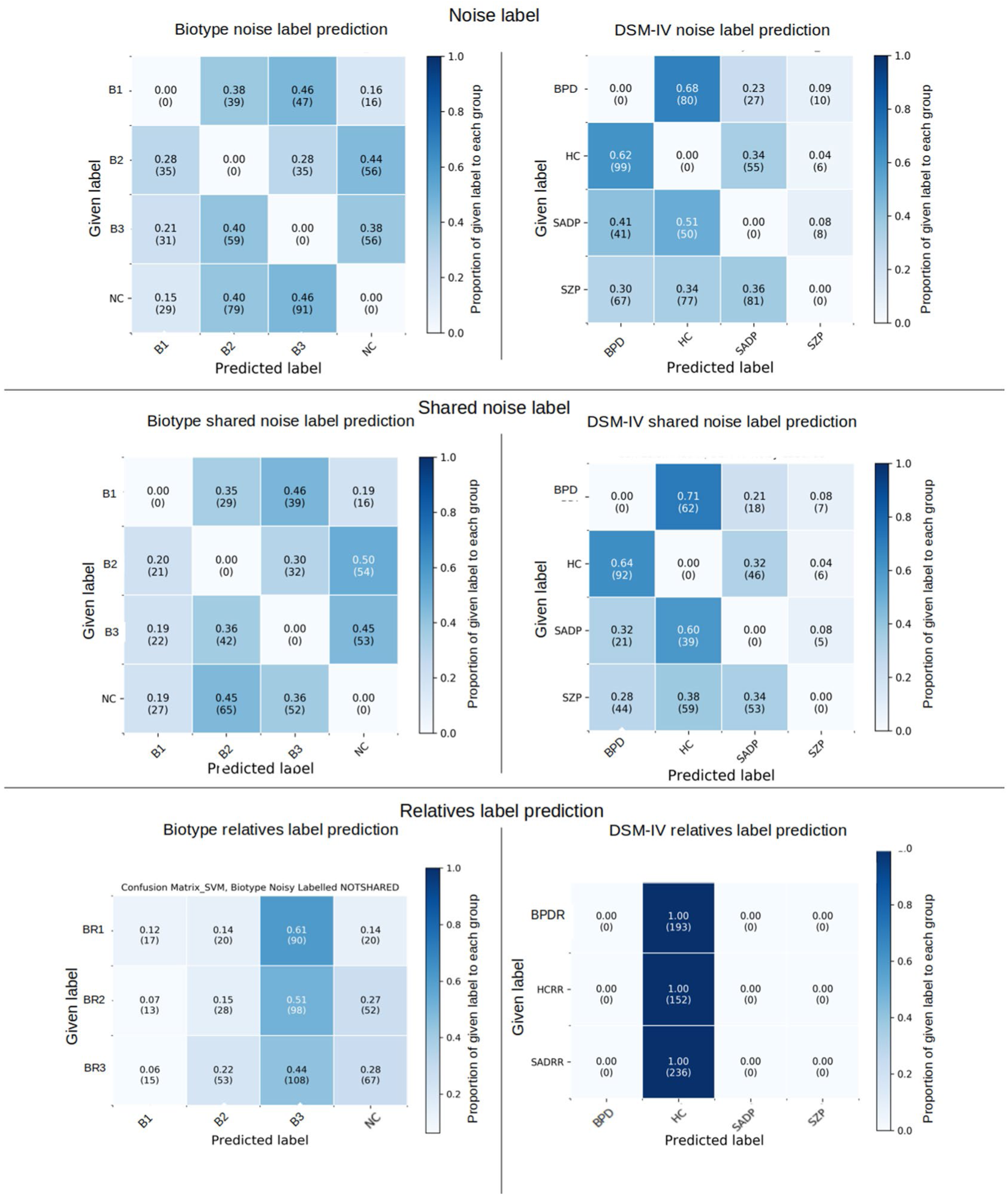
Top: Confusion matrices show label noise subjects related to Biotype or DSM-IV categories separately after convergence to 90% of cleansed data using consensus voting. Middle: Confusion matrices show shared label noise subjects among Biotype and DSM-IV categories after convergence. About 50% of Biotype proband 2 (B2) and 45% of Biotype proband 3 (B3) were relabeled as normal control. Among Biotype proband 1 (B1), 81% of instances identified as noisy label were relabeled as Biotype proband 2 (B2) and Biotype proband 3 (B3). 81% of normal control noisy subjects were also relabeled as Biotype proband 2 (B2) and Biotype proband 3 (B3). In DSM-IV category, 71% of bipolar proband noisy subjects were relabeled as healthy controls, whereas 64% and 32 % of healthy controls were relabeled as bipolar proband and schizoaffective proband respectively. Among schizophrenia label noise subjects, 38% were relabeled as healthy controls and 34% as schizoaffective proband. Bottom: Confusion matrices show the result of predicting labels of relatives using updated labels after the above analysis. Most of the relatives are labeled as Biotype3 (B3) and normal controls in the Biotype categories or as healthy controls in the DSM-IV category.

Among the evaluated criteria, the label noise found in DSM-IV showed more irregularity than Biotype criteria after iterative classification filtering. Among 912 subjects in different groups of DSM-IV, we found that the schizophrenia group showed the most differences. By comparing given labels and new assigned labels, the proportion of schizophrenia group reduced from 26% to 4% of total subjects after convergence and only 6% of schizophrenia subjects remained in the same category after filtering and relabeling. About 62% of schizophrenia subjects were assigned new labels in schizoaffective and bipolar disorder categories and 32% of them were relabeled as healthy controls. Also, from 912 subjects, about 6% of the bipolar group, 2% of healthy control and 6% of the schizoaffective group, were relabeled as schizophrenia. Among different groups of patients in DSM-IV, 50% of bipolar, 20% of schizoaffective and 42% of schizophrenia control were relabeled as healthy control. 56% of healthy controls remained in their group and from remaining healthy control noisy subjects, 27% of them categorized as bipolar, 15% relabeled as schizoaffective and 2% were relabeled as schizophrenia.

On the other hand, for the Biotype groups, about 32% of subjects remained in their original Biotype patient groups and 45% of normal controls remained in their group. Normal controls and Biotype category 3 (B3) changed subjects between each other more than any other two groups in the Biotype criterion. This may because Biotype categories are more based on neuromarkers than clinical symptoms and Biotype category 3 is biologically most similar to normal controls [22]. For the Biotype cleansed relabeled dataset, 51% of relatives were categorized as Biotype B3, 24% of them were labeled as normal control, 17% of them were categorized as Biotype B2 and 8% were categorized as Biotype B1.

T-SNE 2D-projection of original and new labels shows how well subjects categorized based on their labels. First, 100 features were extracted from nonzero voxels using the univariate feature selection method. Next, these features were projected in the 2D plot using tSNE. **Fig 4** shows the t-SNE 2D-projection of the original dataset with label noise in the left panel, and projection using the new suggested labels in the right panel.

**Figure 4.**
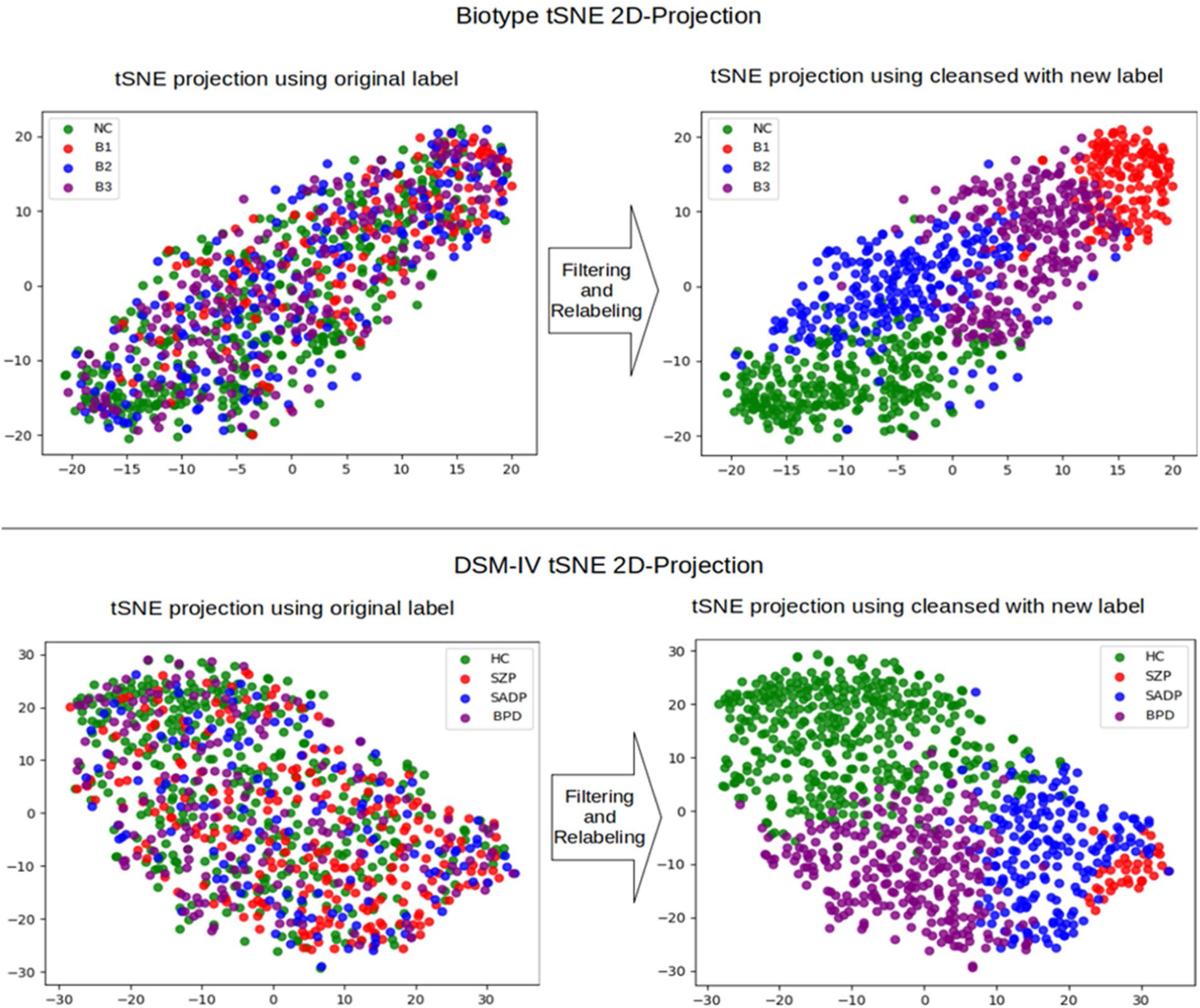
Left panel shows t-SNE 2D-projection of original dataset with label noise and right panel shows 2D-projection using new suggested labels. Top: 2D-projection using Biotype labels. Bottom: 2D-projetion using DSM-IV labels. In both categories of Biotype and DSM-IV, affinities between data points shows subjects in different group of disease and healthy control overlap in noisy data. After identifying label noise and relabeling, the similarity between data points is more obvious and subjects with the same labels are close together. Embedded data into 2D space using original labels does not support the fact that subjects labeled based on their similarities. It is hard to interpret how groups differentiate from each other using original label, because there is considerable overlap between subjects. Cleansed dataset with new suggested label shows there is gradient pattern in both DSM-IV and Biotype categories from healthy/normal control to most severe cases of Biotype B1 or schizophrenia probands contain other mild cases in between.

Voxel-wise t-tests were run on non-zero voxels between each category. Then FDR correction was applied to correct for multiple comparisons. Voxels showing significant differences at a significance level α=0.05 were identified. **Fig 5, 6, 7, 8** show the results of statistical testing between each group. **Fig 5, 6** show gray matter brain maps results using Biotype cleansed labels and Biotype original given label (old label) containing noise. **Fig 7, 8** show gray matter brain maps results using DSM-IV cleansed labels and DSM-IV original diagnostic labels. Results who more voxels showing significant differences when using the cleansed data, and more voxels showing significant differences in the DSM compared to the Biotype data. Importantly the regions that are shown are consistent with previous work [10] [11] (for example in schizophrenia we find reductions in bilateral temporal and insula regions as well as medial frontal).

**Figure 5.**
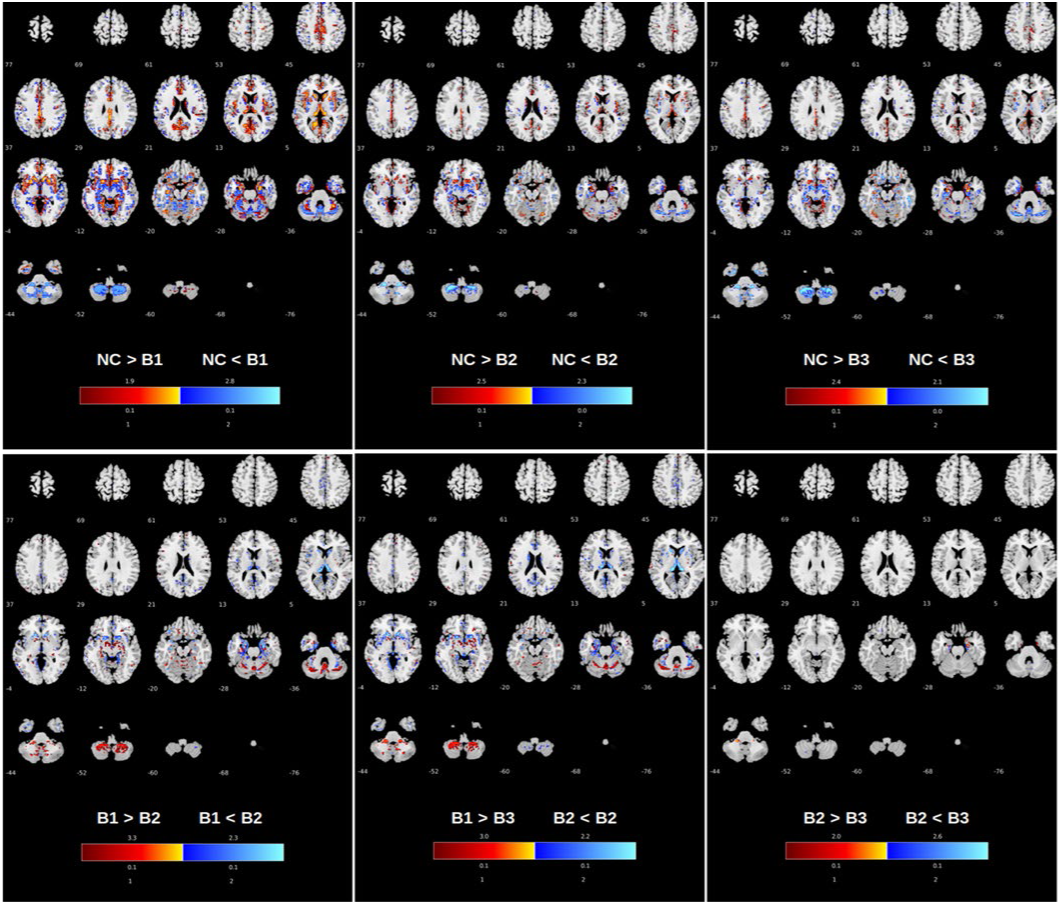
Gray matter map results of voxel wise t-tests between 4 Biotype groups after data cleansing using classification voting filtering. Top row) NC vs. B1, NC vs. B2, NC vs. B3. Bottom row) B1 vs. B2, B1 vs. B2, B2 vs. B3. Gray matter contrast between normal controls and Biotype probands shows they have differences with different levels in overlapped regions after cleaning and relabeling individuals. Gray matter differences between normal control and Biotype proband 1 (B1) have strongest separation among the other comparison tests. The gray matter contrast is lower between healthy controls and Biotype proband 3 (B3). Biotype proband 2 (B2) and Biotype proband 3 (B3) show the fewest differences.

**Figure 6.**
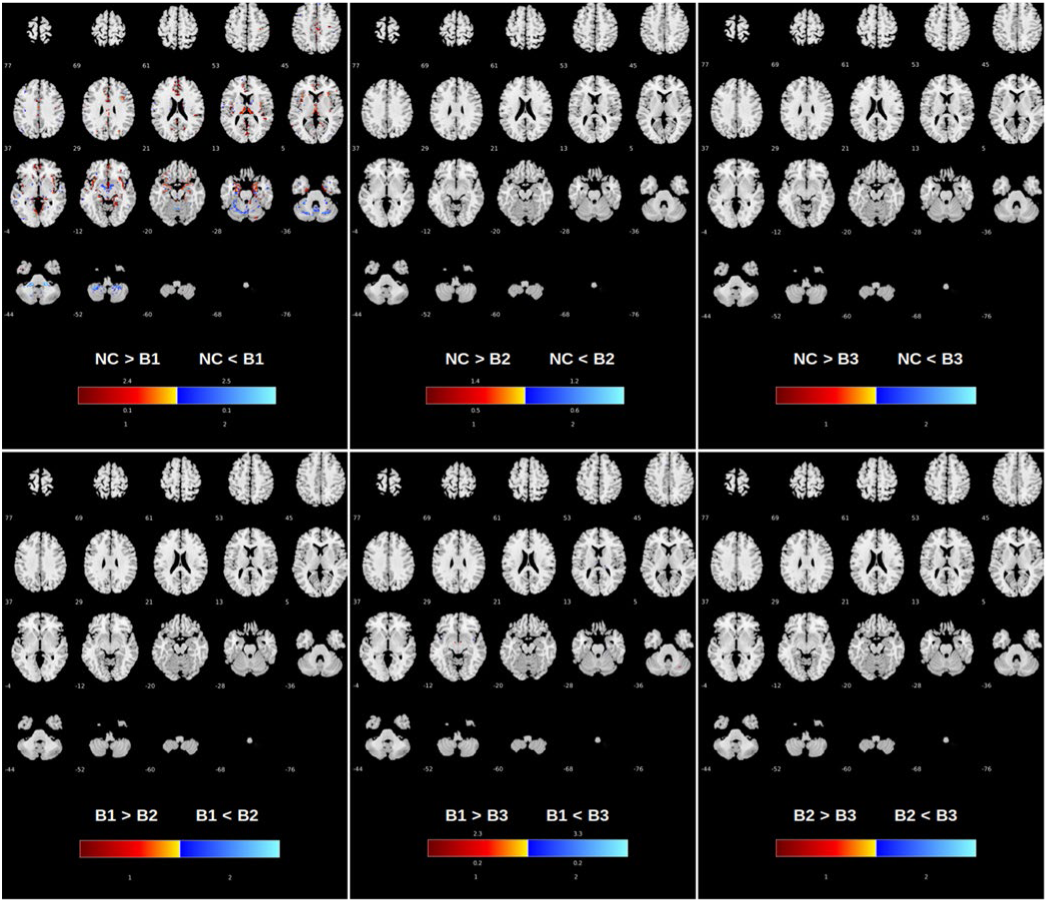
Gray matter map results of voxel wise t-tests between 4 Biotype groups on given labels. Top row: NC vs. B1, NC vs. B2, NC vs. B3; Bottom row: B1 vs. B2, B1 vs. B2, B2 vs. B3. Gray matter contrast between normal controls and Biotype probands only shows significant difference in some regions between normal controls and Biotype proband 1 (B1) using original labels. Gray matter density group differences between other groups do not show differences.

**Figure 7.**
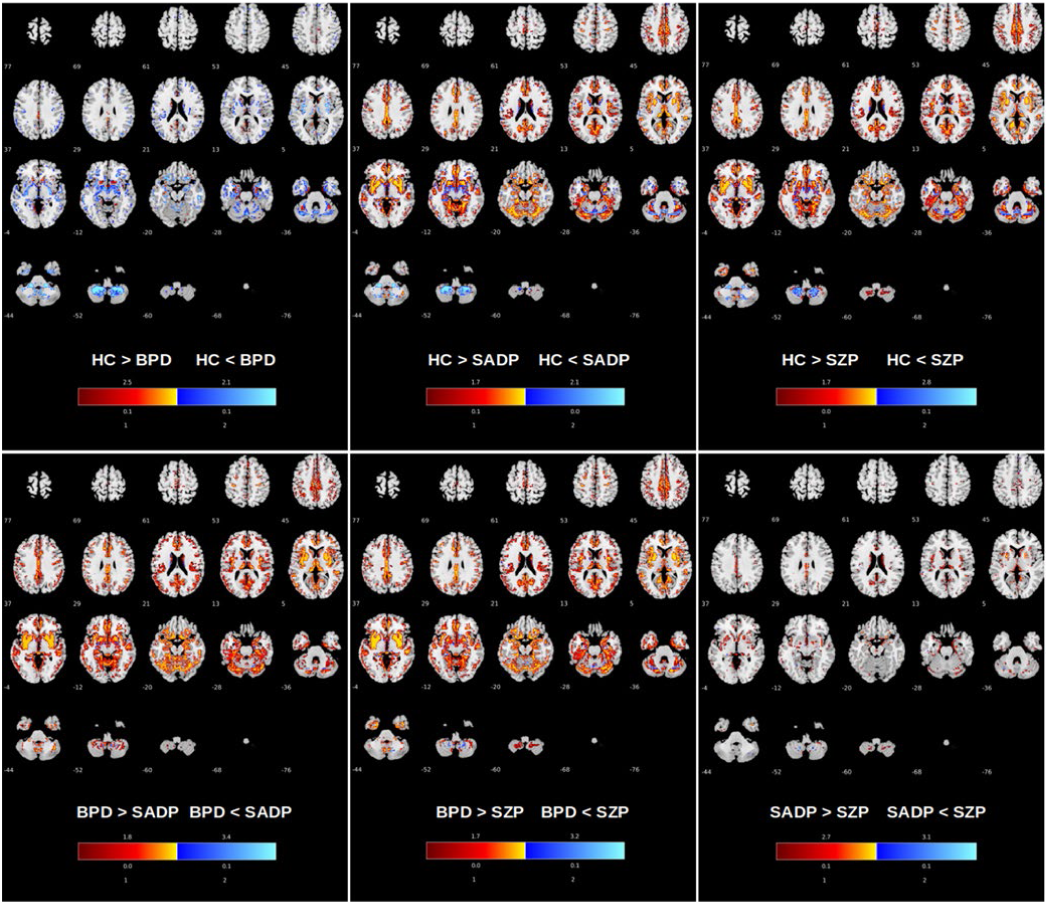
Gray matter map results of voxel wise t-tests between 4 DSM-IV groups after data cleansing using classification voting filtering. Top row) BPD vs. HC, HC vs. SADP, HC vs. SZP. Bottom row) BPD vs. SADP, BPD vs. SZP, SADP vs. SZP. Gray matter contrast between healthy controls and DSM-IV probands shows they have differences with different levels in some overlapped regions after cleaning and relabeling individuals. Gray matter difference between healthy control and bipolar proband (BPD) has strongest separation among the other. The gray matter contrast shows differences between healthy control and schizophrenia and schizoaffective probands as well. Also, group differences could be observed between the bipolar proband group vs. schizophrenia and bipolar proband group vs. schizoaffective proband groups. However, schizophrenia and schizoaffective groups did not separate after data cleansing and relabeling.

**Figure 8.**
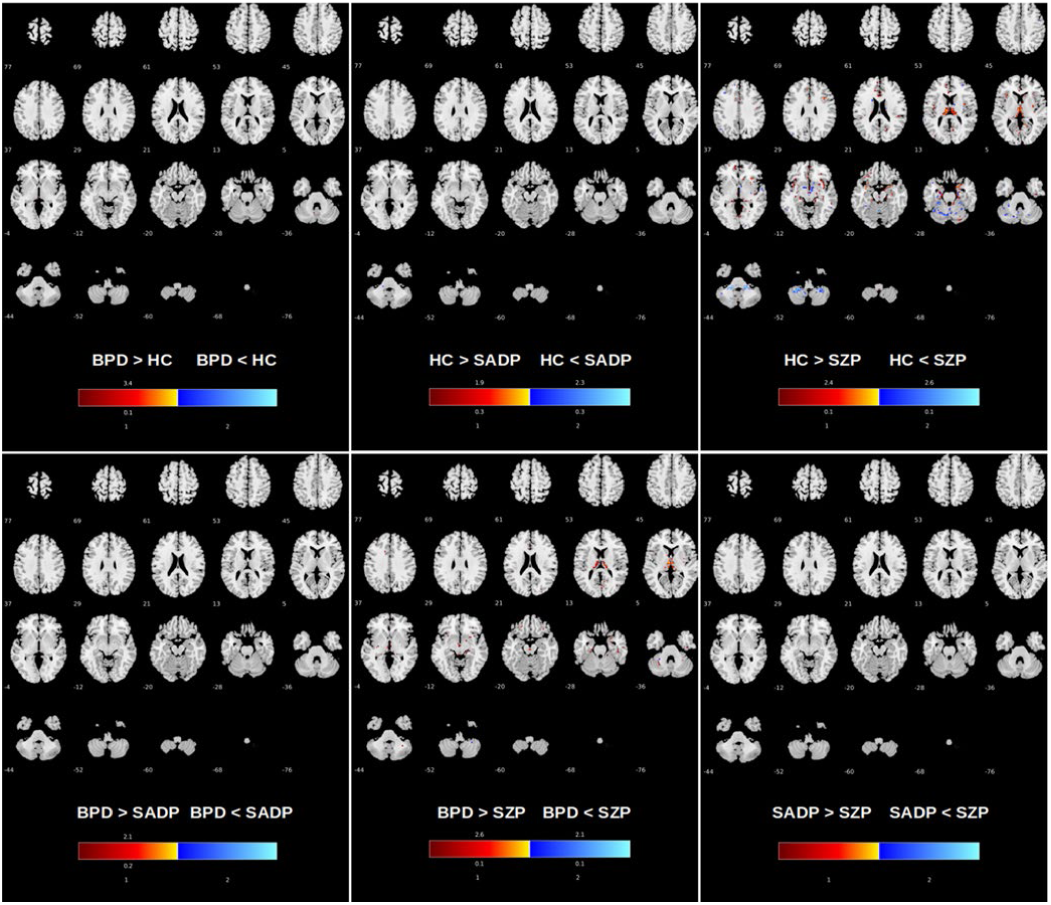
Gray matter map results of voxel wise t-tests between 4 DSM-IV groups on given labels. Top row: BPD vs. HC, HC vs. SADP, HC vs. SZP. Bottom row) BPD vs. SADP, BPD vs. SZP, SADP vs. SZP. Gray matter contrast was only significant in some areas between healthy controls and schizophrenia proband using the original labels. Group differences between other groups were not found.

**Table 3** shows regions where the statistical tests are significantly different after FDR multiple comparisons correction on the p-values. When using the original dataset, we did not find differences in as many regions. There were significant differences in some regions between healthy controls and schizophrenia and bipolar probands and schizophrenia. Significantly different brain regions for the relabeled data are indicated with ● and by ◊ in the original dataset.

**Table 3.**
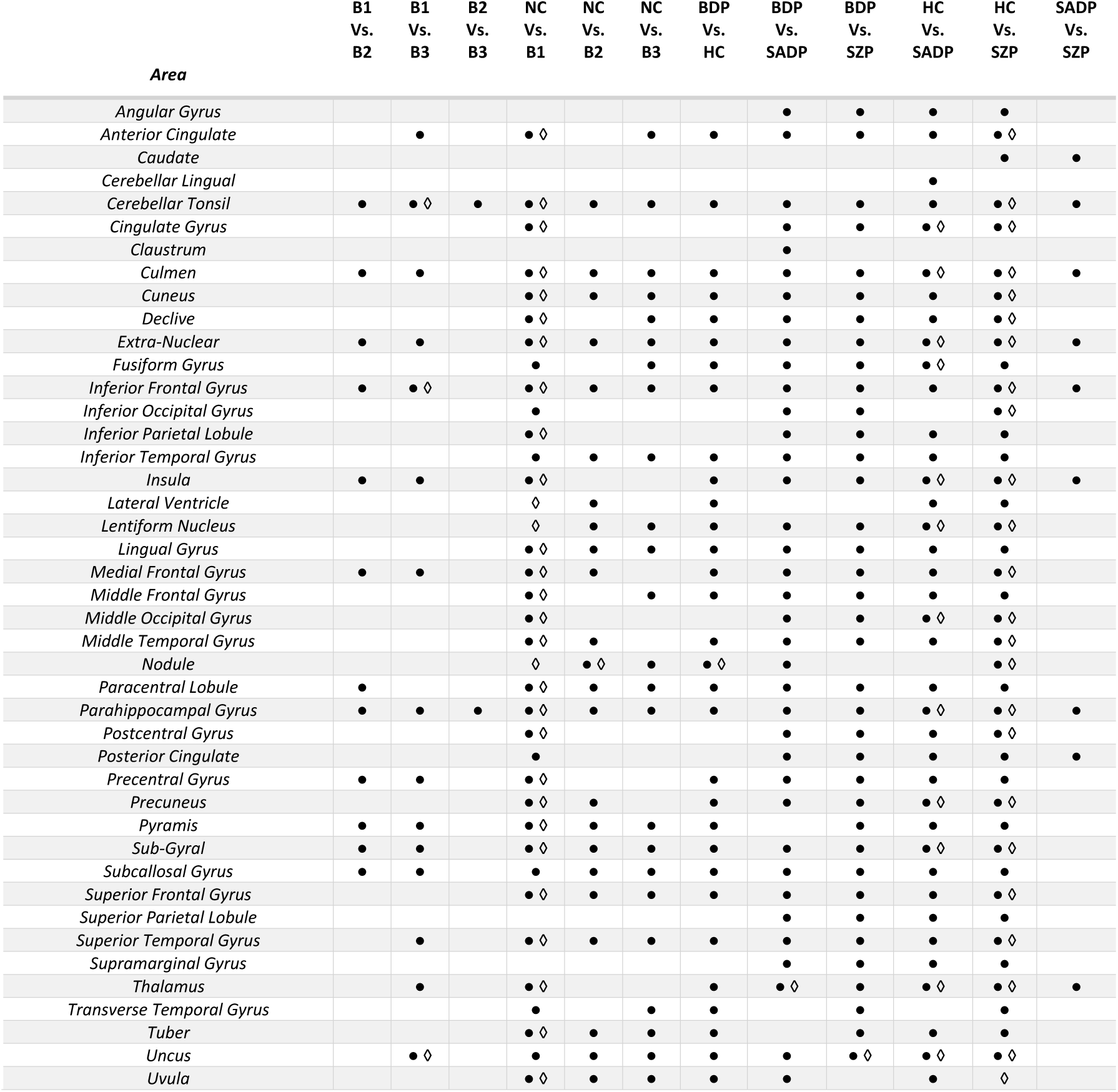
Area of containing voxels obtained by statistical significant test on the relabeled data ● and original data ◊

## 4 DISCUSSION

Current diagnostic categories for mental illness are not biologically based and exhibit considerable heterogeneity within diagnosis and overlap across diagnoses [29, 30]. Most imaging biomarker studies do not surprisingly use the DSM categories as the ground truth, which, while informative, does not help us move beyond the known validity issues. In this work, we present a first towards addressing the known problems with existing labeling schemes but providing a way for them to be updated by biological data (e.g. brain imaging). The approach we use uses a current categorization approach as input but then updates are using additional biological data by assuming there is noise in the assignment process. We show that the MRI data is pushing against the labeling assigned via two different diagnostic categorization approaches (DSM & biotypes). Then using existing categories, we ask how the data should be categorized within each scheme to reduce the label noise. We are currently exploring extending this work via semi-supervised approaches and to nominate new categories based on the biological data alone. This is an extremely difficult task which has been underexplored in the context of brain imaging data and we hope to contribute more on this going forward.

Many studies have been done to address label noise issues in different domains such as medical imaging [31, 32, 33, 34, 35, 36]. However, label noise is still an open question in computer-aided diagnosis systems. It is arguably made worse by the use of advanced approaches like deep learning, which also typically require a ground truth [32]. The problem is very difficult for mood and psychosis disorder, which share symptoms. It is often challenging to obtain reliable labels for the classification task and in reality, because of many reasons such as insufficient information or even expert’s mistakes or poor data quality [1] we have to deal with labels that are polluted by label noises [2].

Additionally, data acquisition, quality control and preprocessing steps can have an impact on the estimated label noise, though we do careful quality control in order to mitigate these aspects as much as possible. Mislabeled instances may be included in any proportion of data i.e., training, validation and testing dataset which affect the performance of model evaluation and produce unreliable results. Also, there is often considerable overlap among the subjects and no clear boundaries between the groups. For all these reasons and a lack of ground truth in neuroimaging studies, identification of label noise becomes extremely difficult.

Voting filtering is one approach that has been proposed to try to identify label noise and involves removing an instance when all learners agree [1] [34] [37]. We approach this from a data cleansing perspective, that is, we first identify the noisy labels. One advantage of a data cleansing approach is it reduces the complexity of the model. It also increases the performance of the classifier; however, as we saw, it may remove a large number of cases. Indeed, if those cases are truly noisy (either because the label or the labeling system is inaccurate or incomplete), then they should be excluded. To avoid overcleansing problems and mitigating removing minority instances using classification filtering in imbalanced datasets, we used a subsampling approach. We used a random under-sampling approach to handle imbalanced classes and resampled the majority class randomly and uniformly. This was done repeatedly till all the instances of the majority classes were visited at least one time for each cross-validation iteration. Future studies will explore the other possible approaches for addressing imbalanced data.

Simulation study (described in supplemental) supports our view that one of the consequences of label noise is that it decreases the performance and accuracy of the model. The presence of label noise can disrupt the underlying patterns we are trying to discover. Our analysis of real data also proves this fact that label noise blurred the underlying patterns. The 2D projection using original labels shows lack of similarities between each subject in the 2D space within groups. This does not support the fact that there is a relationship between probands belonging to the same group. However, the cleansed 2D projection shows there is a gradient pattern between groups from healthy controls to the most severe Biotype case B1 or from healthy controls to schizophrenia for DSC. Results were promising, but additional work is needed. For example, some of the results suggest a gradient pattern between groups from healthy controls to more several patients. Thus the use of additional subcategories or using dimensional information (as suggested in the NIMH RDoC approach)instead of a categorical approach may provide a more meaningful representation of relabeled subjects and underlying relation between them. More detailed mood disorder diagnoses will lead to more effective treatments [38]

Cleansed data showed many more significant voxels than did the original data. In addition, the DSM-IV cleansed data showed more significant voxels than the Biotype cleansed data. Interestingly, the cleansed dataset showed many more significant differences, providing possible support for our approach. Reclassified relatives had the same distinct features in brain regions more similar to DSM-IV healthy control and Biotype B3 and Biotype normal control in the cleansed dataset.

Reclassifying subjects does not suggest a patient is not actually sick or a healthy person had mental disorders. Relabeling just shows how the subjects can be grouped in more homogeneous categories based on the sMRI features. However, it shows very clearly that categories are not reflecting the underlying biology (as measures by sMRI) well. In addition, the fact that 1) the relabeled data showed a clear gradient from the most to least severe categories, and 2) there were more voxelwise group differences in regions that were consistent with what might be expected, in the cleansed data, provides intriguing evidence and supports continued work in this direction.

Also, the main reason for choosing unsmoothed images was to avoid removing relevant information. Indeed, if we smooth the images, the classification results are lower. Future work can include incorporating multiple types of data (e.g. EEG, structural MRI and functional MRI). In addition, allowing the approach to developing new categories (e.g. via splitting and merging) is another interesting topic for future work. Another interesting and important avenue of study is to investigate the individuals who were identified as having noisy labels in more depth (i.e. the boundary cases). We are planning to do this in future work. Ultimately the results need to be validated clinically.

## 5 CONCLUSION

In this paper, we proposed a novel approach to estimate label noise prevalent due to incorrect diagnostic classification. We used iterative classification voting filtering using an SVM model. We applied our method to brain imaging in the context of a multi-label imbalanced data from various psychosis disorders and healthy controls. Overall accuracy increases and converges after iterating classification filtering and relabeling steps. The proposed method provides a promising approach for feature extraction of brain images even on noisy datasets by assigning new labels to those inconsistently diagnosed/labeled individuals over multiple iterations. This method identified noisy label samples and suggested new labels for them with the current noisy dataset without requiring extra clean samples in any classification domain by estimating appropriate models and finding optimal hyper-parameters for each classification task. Our method shows that although there was a similar proportion of label noise in multiclass Biotype and DSM-IV categories, label noise distributions were more irregular in DSM-IV than in Biotype categories; however, DSM-IV data showed more significant voxels when evaluating group differences on cleansed data. Our method showed a transparent gradient from the most to least clinically severe groups. Our hope is that this represents an initial step towards a semi-blind categorization approach that is both informed by clinical as well as high-dimensional biological data.

## 6 ACKNOWLEDGMENTS

The research reported in this work was supported by the National Institute of Mental Health under award numbers R01EB005846 and 1R01MH104680.

## REFERENCES

[1] B. Frenay and M. Verleysen, “Classification in the presence of label noise: a survey,” IEEE transactions on neural networks and learning systems, vol. 25, no. 5, pp. 845–869, 2013.

[2] I. Bross, “Misclassification in 2 × 2 tables,” Biometrics, vol. 10, no. 4, pp. 478–486, 1954.

[3] A. Hadgu, “The discrepancy in discrepant analysis,” The Lancet, vol. 348, no. 9027, pp. 592–593, 1996.

[4] B. Frenay, A. Kaban and others, A comprehensive introduction to label noise, 2014.

[5] A. Abrol, H. Rokham and V. D. Calhoun, “Diagnostic and Prognostic Classification of Brain Disorders Using Residual Learning on Structural MRI Data,” In 41st Annual International Conference of the IEEE Engineering in Medicine & Biology Society (EMBC), Berlin, 2019.

[6] Z. Wang, S. A. Meda, M. S. Keshavan, C. A. Tamminga, J. A. Sweeney, B. A. Clementz, D. J. Schretlen, V. D. Calhoun, S. Lui and G. D. Pearlson, “Large-scale fusion of gray matter and resting-state functional MRI reveals common and distinct biological markers across the psychosis spectrum in the B-SNIP cohort,” Frontiers in psychiatry, vol. 6, p. 174, 2015.

[7] G. D. Pearlson, B. A. Clementz, J. A. Sweeney, M. S. Keshavan and C. A. Tamminga, “Does biology transcend the symptom-based boundaries of psychosis?,” Psychiatric Clinics, vol. 39, no. 2, pp. 165–174, 2016.

[8] T. R. Insel and B. N. Cuthbert, “Brain disorders? Precisely,” Science, vol. 348, no. 6234, pp. 499–500, 2015.

[9] C. P. Langlotz, B. Allen, B. J. Erickson, J. Kalpathy-Cramer, K. Bigelow, T. S. Cook, A. E. Flanders, M. P. Lungren, D. S. Mendelson, J. D. Rudie, G. Wang and K. Kandarpa, “A roadmap for foundational research on artificial intelligence in medical imaging: From the 2018 NIH/RSNA/ACR/The Academy Workshop,” Radiology, vol. 291, no. 3, pp. 781–791, 2019.

[10] B. L. Amann, E. Canales-Rodr{\’\i}guez, M. Madre, J. Radua, G. Monte, S. Alonso-Lana, R. Landin-Romero, A. Moreno-Alc{\’a}zar, C. Bonnin, S. Sarr{\’o}, O.-G. J G. JJ M. N, F.-C. P, G. JM, B. J, S. R and others, “Brain structural changes in schizoaffective disorder compared to schizophrenia and bipolar disorder,” Acta Psychiatrica Scandinavica, vol. 133, no. 1, pp. 23–33, 2016.

[11] H. G. Schnack, M. Nieuwenhuis, N. E. van Haren, L. Abramovic, T. W. Scheewe, R. M. Brouwer, H. E. H. Pol and R. S. Kahn, “Can structural MRI aid in clinical classification? A machine learning study in two independent samples of patients with schizophrenia, bipolar disorder and healthy subjects,” Neuroimage, vol. 84, pp. 299–306, 2014.

[12] H. Falakshahi, V. M. Vergara, J. Liu, D. H. Mathalon, J. M. Ford, J. Voyvodi, B. Mueller, A. Belger, S. McEwen, S. Potkin, A. Preda, H. Rokham, J. Sui, J. A. Turner, S. Plis and V. D. Calhoun, “Meta-modal Information Flow: A Method for Capturing Multimodal Modular Disconnectivity in Schizophrenia,” IEEE Transactions on Biomedical Engineering, 2020.

[13] H. Rokham, H. Falakshahi and V. D. Calhoun, “A data-driven approach for stratifying psychotic and mood disorders subjects using structural magnitude resonance imaging data,” In Medical Imaging 2020: Computer-Aided Diagnosis, 2020.

[14] R. J. Hickey, “Noise modelling and evaluating learning from examples,” Artificial Intelligence, vol. 82, no. 1–2, pp. 157–179, 1996.

[15] J. R. Quinlan, “Induction of decision trees,” Machine learning, vol. 1, no. 1, pp. 81–106, 1986.

[16] X. Zhu and X. Wu, “Class noise vs. attribute noise: A quantitative study,” {Artificial intelligence review, vol. 22, no. 3, pp. 177–210, 2004.

[17] D. Rolnick, A. Veit, S. Belongie and N. Shavit, “Deep learning is robust to massive label noise,” arXiv preprint 1705.10694, 2017.

[18] C. Sun, A. Shrivastava, S. Singh and A. Gupta, “Revisiting unreasonable effectiveness of data in deep learning era,” In Proceedings of the IEEE international conference on computer vision, 2017.

[19] C. A. Tamminga, G. Pearlson, M. Keshavan, J. Sweeney, B. Clementz and G. Thaker, “Bipolar and schizophrenia network for intermediate phenotypes: outcomes across the psychosis continuum,” Schizophrenia bulletin, vol. 40, no. Suppl_2, pp. S131--S137, 2014.

[20] C. A. Tamminga, E. I. Ivleva, M. S. Keshavan, G. D. Pearlson, B. A. Clementz, B. Witte, D. W. Morris, J. Bishop, G. K. Thaker and J. A. Sweeney, “Clinical phenotypes of psychosis in the Bipolar-Schizophrenia Network on Intermediate Phenotypes (B-SNIP),” American Journal of psychiatry, vol. 170, no. 11, pp. 1263–1274, 2013.

[21] E. I. Ivleva, A. S. Bidesi, M. S. Keshavan, G. D. Pearlson, S. A. Meda, D. Dodig, A. F. Moates, H. Lu, A. N. Francis, N. Tandon and others, “Gray matter volume as an intermediate phenotype for psychosis: Bipolar-Schizophrenia Network on Intermediate Phenotypes (B-SNIP,” American Journal of Psychiatry, vol. 170, no. 11, pp. 1285–1296, 2013.

[22] E. I. Ivleva, B. A. Clementz, A. M. Dutcher, S. J. Arnold, H. Jeon-Slaughter, S. Aslan, B. Witte, G. Poudyal, H. Lu, S. A. Meda and others, “Brain structure biomarkers in the psychosis biotypes: findings from the bipolar-schizophrenia network for intermediate phenotypes,” Biological psychiatry, vol. 82, no. 1, pp. 26–39, 2017.

[23] C. R. Jack Jr, M. A. Bernstein, N. C. Fox, P. Thompson, G. Alexander, D. Harvey, B. Borowski, P. J. Britson, J. L. Whitwell, C. Ward and others, “The Alzheimer’s disease neuroimaging initiative (ADNI): MRI methods,” Journal of Magnetic Resonance Imaging: An Official Journal of the International Society for Magnetic Resonance in Medicine, vol. 27, no. 4, pp. 685–691, 2008.

[24] J. Ashburner and K. J. Friston, “Unified segmentation,” Neuroimage, vol. 26, no. 3, pp. 839–851, 2005.

[25] F. Pedregosa, G. Varoquaux, A. Gramfort, V. Michel, B. Thirion, O. Grisel, M. Blondel, P. Prettenhofer, R. Weiss, V. Dubourg, J. Vanderplas, A. Passos, D. Cournapeau, M. Brucher, M. Perrot and E. Duchesnay, “Scikit-learn: Machine Learning in {P}ython,” Journal of Machine Learning Research, vol. 12, pp. 2825–2830, 2011.

[26] B. Mwangi, T. S. Tian and J. C. Soares, “A review of feature reduction techniques in neuroimaging,” Neuroinformatics, vol. 12, no. 2, pp. 229–244, 2014.

[27] R. E. Bellman, Adaptive control processes: a guided tour, vol. 2045, Princeton university press, 2015.

[28] L. v. d. Maaten and G. Hinton, “Visualizing data using t-SNE,” Journal of machine learning research, vol. 9, pp. 2579–2605, 2008.

[29] K. Allsopp, J. Read, R. Corcoran and P. Kinderman, “Heterogeneity in psychiatric diagnostic classification,” Psychiatry research, vol. 279, pp. 15–22, 2019.

[30] C. M. Olbert, G. J. Gala and L. A. Tupler, “Quantifying heterogeneity attributable to polythetic diagnostic criteria: theoretical framework and empirical application.,” Journal of Abnormal Psychology, vol. 123, no. 2, p. 452, 2014.

[31] E. Calli, E. Sogancioglu, E. T. Scholten, K. Murphy and B. van Ginneken, “Handling label noise through model confidence and uncertainty: application to chest radiograph classification,” In Medical Imaging 2019: Computer-Aided Diagnosis, 2019.

[32] C. Xue, Q. Dou, X. Shi, H. Chen and P. A. Heng, “Robust Learning at Noisy Labeled Medical Images: Applied to Skin Lesion Classification,” arXiv preprint 1901.07759, 2019.

[33] M. Pechenizkiy, A. Tsymbal, S. Puuronen and O. Pechenizkiy, “Class noise and supervised learning in medical domains: The effect of feature extraction,” In 19th IEEE Symposium on Computer-Based Medical Systems (CBMS’06), 2006.

[34] D. Gamberger, N. Lavrac and C. Groselj, “Experiments with noise filtering in a medical domain,” In ICML, 1999, pp. 143–151.

[35] S. Ji and J. Ye, “Generalized linear discriminant analysis: a unified framework and efficient model selection,” In IEEE Transactions on Neural Networks, 2008.

[36] K. Robbins, S. Joseph, W. Zhang, R. Rekaya and J. Bertrand, “Classification of incipient Alzheimer patients using gene expression data: Dealing with potential misdiagnosis,” Online Journal of Bioinformatics, vol. 7, no. 1, pp. 22–31, 2006.

[37] C. E. Brodley and M. A. Friedl, “Identifying mislabeled training data,” Journal of artificial intelligence research, vol. 11, pp. 131–167, 1999.

[38] G. B. Chand, D. B. Dwyer, G. Erus, A. Sotiras, E. Varol, D. Srinivasan, J. Doshi, R. Pomponio, A. Pigoni, P. Dazzan, R. S. Kahn, H. G. Schnack, M. V. Zanetti, E. Meisenzahl, G. F. Busatto and Crespo-Facor, “Two distinct neuroanatomical subtypes of schizophrenia revealed using machine learning,” Brain, no. 2, 02 2020.

[39] N. Japkowicz and S. Stephen, “The class imbalance problem: A systematic study,” Intelligent data analysis, vol. 6, no. 5, pp. 429–449, 2002.

[40] F. Provost, “Machine learning from imbalanced data sets 101,” In Proceedings of the AAAI’2000 workshop on imbalanced data sets, 2000.

