## Supplemental Materials for "Addressing Inaccurate Nosology in Mental Health: A Multi Label Data Cleansing Approach for Detecting Label Noise from Structural Magnetic Resonance Imaging Data in Mood and Psychosis Disorders"

### Supplementary

**Grid search:**

*1.dimension reduction:* We applied grid search consisted of 3 different dimension reduction algorithms, 1) selecting k feature using analysis of variance (ANOVA), which is computing a univariate ANOVA F-value between groups based on their assigned labels and apply this to select the best features among vectors of features (vector of voxels) among instances. 2) Principal component analysis (PCA) which is a statistical procedure that uses an orthogonal transformation to find principal components which are uncorrelated to each other. 3) FastICA which is an efficient algorithm that performs independent component analysis (ICA). In this study we set 100 components for these three approaches.

*2. Hyper-parameters optimization and parameter selection:* Hyper-parameters are parameters within a learning algorithm which should be defined before model training and fitting. In contrast to ad-hoc hyper-parameters setting, hyper-parameter tuning/optimization refers to identifying the optimal set of hyper-parameters which maximizes the performance of the algorithm among the other options. In this work, to find the best hyper-parameters for the training model, we used a grid search among different hyper-parameters space and settings for the SVM model. Grid search for SVM classifier consists of super vector classifiers with 3 different nonlinear kernels (radial basis function, polynomial and sigmoid) as well as a linear kernel. The search was conducted over a kernel coefficient range of (-4, 4) for the nonlinear kernels. Also (0.0001, 1.0) was used as a regularization parameter space for grid search settings. We used nested cross validation by splitting data into two equal portions (50%) for training and testing in inner cross validation and using 5-fold cross validation for outer cross validation.

This approach seeks the hyper-parameter space through cross validation and proposes the best candidate among possible hyper-parameters values.

**B-SNIP dataset:** The mean and standard deviation of the positive and negative syndrome scale (PANSS) for different categories are shown in **Table 1** which also includes the mood status of each category. PANSS are not available for control subjects.

Data were categorized and labeled using two different approaches. This includes both standard clinical diagnosis based on the DSM-IV and also neurobiological heterogeneity defined groups, called Biotypes [1]. Each Biotype category contained individuals with all DSM psychosis categories and vice versa (see **Table S1**). According to [1], "Biotypes" (biologically distinctive phenotypes) refer to neurobiologically distinct subgroups of psychosis cases independent of clinical phenomenology that differentiated people with psychosis from healthy controls. Biotype group B1 manifest cases with impaired cognitive control and poor sensorimotor function, group B2 was characterized by impaired cognitive control but exaggerated sensorimotor response, and group B3 presented near normal cognitive and sensorimotor functions [2]. DSM categories included bipolar disorder (BPD) with psychosis, schizoaffective disorder (SADP), schizophrenia (SZP) and also healthy control (HC) subjects. In this study, we separately evaluated the DSM and Biotype labeled data.

***Table S1*** *DSM-IV diagnostic criteria features of all Biotype groups*

| DSM-IV | BIOTYPE | # OF INDIVIDUALS |
|---|---|---|
| **BPD** | B1 | 25 |
| | B2 | 57 |
| | B3 | 94 |
| **SADP** | B1 | 33 |
| | B2 | 48 |
| | B3 | 53 |
| **SZP** | B1 | 89 |
| | B2 | 80 |
| | B3 | 71 |
| **HC** | NC | 362 |

The B-SNIP dataset also includes 581 patients' relatives (Biotype patient relatives) which consist of 193 bipolar relatives (BPDR), 152 schizoaffective disorder relatives (SADPR) and 236 schizophrenia relatives (SZPR). Within the Biotype categorization, 147 of them are relatives of Biotype group 1 (BR1), 191 subjects are relatives of Biotype group 2 (BR2), 116 subjects are relatives of Biotype group 3 (BR2). Also, among 581 relative subjects only 82 of them have PANSS available. 26 bipolar proband relatives with average PANSS 52.3 +/- 14.5, 28 schizoaffective proband relatives with average PANSS 58.14 +/- 17 and 28 schizophrenia proband relatives with average PANSS 58.28 +/- 17.26.

The medication status of the individuals reported in this dataset is shown in Table 3. About 95% of bipolar disorder subjects, 95% of the schizoaffective proband and 92% of schizophrenia patients reported that they were using psychotropic medications.

***Table S2*** *Medicine status of the subjects in B-SNIP dataset*

| MEDICENE TYPE | REPORTED | TOTAL NUMBERS | BPD | | SADP | | SZP | | HC | |
|---|---|---|---|---|---|---|---|---|---|---|
| | | | *N* | *%* | *N* | *%* | *N* | *%* | *N* | *%* |
| **ANY PSYCHOTROPIC MEDICATIONS** | Unknown | 18 | 2/176 | 1.14% | | | 4/240 | 1.67% | 12/362 | 3.31% |
| | NO | 370 | 13/176 | 7.39% | 7/134 | 5.22% | 13/240 | 5.42% | 337/362 | 93.09% |
| | YES | 524 | 161/176 | 91.48% | 127/134 | 94.78% | 223/240 | 92.92% | 13/362 | 3.59% |
| **ANTIPSYCHOTIC** | Unknown | 18 | 2/176 | 1.14% | | | 4/240 | 1.67% | 12/362 | 3.31% |
| | NO | 430 | 44/176 | 25.00% | 16/134 | 11.94% | 20/240 | 8.33% | 350/362 | 96.69% |
| | YES | 464 | 130/176 | 73.86% | 118/134 | 88.06% | 216/240 | 90.00% | | |
| **ANTIDEPRESSANT** | Unknown | 18 | 2/176 | 1.14% | | | 4/240 | 1.67% | 12/362 | 3.31% |
| | NO | 652 | 98/176 | 55.68% | 60/134 | 44.78% | 150/240 | 62.50% | 344/362 | 95.03% |
| | YES | 242 | 76/176 | 43.18% | 74/134 | 55.22% | 86/240 | 35.83% | 6/362 | 1.66% |
| **MOOD STABILIZER** | Unknown | 18 | 2/176 | 1.14% | | | 4/240 | 1.67% | 12/362 | 3.31% |
| | NO | 639 | 51/176 | 28.98% | 57/134 | 42.54% | 181/240 | 75.42% | 350/362 | 96.69% |
| | YES | 255 | 123/176 | 69.89% | 77/134 | 57.46% | 55/240 | 22.92% | | |
| **ANXIOLYTIC SEDATIVES HYPNOTIC** | Unknown | 18 | 2/176 | 1.14% | | | 4/240 | 1.67% | 12/362 | 3.31% |
| | NO | 741 | 120/176 | 68.18% | 97/134 | 72.39% | 181/240 | 75.42% | 343/362 | 94.75% |
| | YES | 153 | 54/176 | 30.68% | 37/134 | 27.61% | 55/240 | 22.92% | 7/362 | 1.93% |
| **ANTICHOLINERGIC ANTIPARKINSONIAN** | Unknown | 18 | 2/176 | 1.14% | | | 4/240 | 1.67% | 12/362 | 3.31% |
| | NO | 819 | 158/176 | 89.77% | 114/134 | 85.07% | 197/240 | 82.08% | 350/362 | 96.69% |
| | YES | 75 | 16/176 | 9.09% | 20/134 | 14.93% | 39/240 | 16.25% | | |
| **MISCELLANEOUS, PSYCHOTROPIC CENTRALLY ACTIVE** | Unknown | 18 | 2/176 | 1.14% | | | 4/240 | 1.67% | 12/362 | 3.31% |
| | NO | 880 | 170/176 | 96.59% | 129/134 | 96.27% | 231/240 | 96.25% | 350/362 | 96.69% |
| | YES | 14 | 4/176 | 2.27% | 5/134 | 3.73% | 5/240 | 2.08% | | |
| **STIMULANTS** | Unknown | 18 | 2/176 | 1.14% | | | 4/240 | 1.67% | 12/362 | 3.31% |
| | NO | 855 | 156/176 | 88.64% | 127/134 | 94.78% | 223/240 | 92.92% | 349/362 | 96.41% |
| | YES | 39 | 18/176 | 10.23% | 7/134 | 5.22% | 13/240 | 5.42% | 1/362 | 0.28% |

Using the Alzheimer's Disease Neuroimaging Initiative protocol (http://adni.loni.usc.edu/methods/documents/mri-protocols/), and appropriate for scanner brands, T1-weighted structural images were acquired on 3T magnets across different sites; T1-weighted magnetization prepared rapid acquisition gradient-echo or inversion recovery-prepared spoiled gradient recoil sequences [2]. **Table S3** shows the parameter settings used for the scanners.

*Table S3* *B-SNIP dataset different parameter settings for the scanners*

| *Scanner* | | *shot interval* | *inversion time* | *TR* | *TE* | *flip angle* | *FOV* | *matrix* | *in-plane resolution* | *slices* | *slice thickness* | *voxel size* | *total scan duration* |
|---|---|---|---|---|---|---|---|---|---|---|---|---|---|
| *GE Signa* | sagittal slab | 2300 ms | 700 ms | 6.99 ms | 2.85 ms | 8º | 260×260 mm$^2$ | 256×256 | 1×1 mm$^2$ | 166 | 1.2mm | 1×1×1 0.2mm$^3$ | 10min 28sec |
| *Philips Achieva* | sagittal slab | 3000 ms | 846 ms | 6.8ms | 3.1ms | 8º | 256×240 mm$^2$ | 256×240 | 1×1 mm$^2$ | 170 | 1.2mm | 1×1×1 0.2mm$^3$ | 9min 19sec |
| *Siemens Allegra* | sagittal slab | 2300 ms | 900 ms | 7.2ms | 2.91 ms | 9º | 256×240 mm$^2$ | 256×240 | 1×1 mm$^2$ | 160 | 1.2mm | 1×1×1 0.2mm$^3$ | 9min 14sec |
| *Siemens Trio* | sagittal slab | 2300 ms | 900 ms | 6.8ms | 2.91 ms | 9º | 256×240 mm2 | 256×240 | 1×1 mm$^2$ | 160 | 1.2mm | 1×1×1 0.2mm$^3$ | 9min 14sec |
| *GE Signa HDxt* | sagittal slab | 2300 ms | 650 ms | 7.0ms | 3.0ms | 8º | 256×256 mm$^2$ | 176x256x176 | 1×1 mm$^2$ | 166 | 1.2mm | 1×1×1 0.2mm3 | 10min9sec |
| *Siemens Trio* | sagittal slab | 2300 ms | 900 ms | 6.8ms | 2.74 ms | 8º | 176×256 mm$^2$ | 176x256x176 | 1×1 mm$^2$ | 160 | 1.2mm | 1×1×1 0.2mm$^3$ | 10min9sec |

**Biotype:** Biotypes leverage neurobiological heterogeneity among psychosis cases in combination with information derived by laboratory tasks that assess brain function at the neurocognitive/perceptual level[1]. By integrating different biomarkers e.g. neuropsychological, stop signal, saccadic control, and auditory stimulation paradigms and other data external validating measures like social functioning, EEG, family biomarkers, and clinical information, nine variables were derived to capture neurobiological variance among the different psychosis cases which leads to three neurobiologically distinct psychosis biotypes which were distinctive and did not map to clinical boundaries. Note the structural MRI data were not used to derive the biotypes used in this study.

Probands and their first relatives assessed with the structured clinical interview for DSM-IV to evaluate psychosis spectrum personality traits. Healthy subjects and their first-degree relatives did not have history of psychotic or bipolar disorder records. The majority of probands and a minor subset of relatives were taking psychotropic medications [1].

For laboratory paradigms, there were multiple such variables, five saccade variables two stop signal variables and 31 EEG variables. PCA data reduction applied for each paradigm set (saccades, stop signal task, EEG) which leads to two saccades, one stop signal, and five EEG components. Then using k-means clustering for class formation, 3 subgroups (biotypes) were used to capture cognitive-perceptual classification variance among the participants with psychosis [1].

**Simulation:** To validate our approach, we conducted a simulation on handwritten digit images. We included images of digits 1, 3, 5, 8 because their shapes' similarity makes the prediction more challenging. The images have dimension 64 pixels (8×8) **Fig S1**. We selected and shuffled truth labels of randomly chosen digit images of different proportions in the dataset. **Fig S2** shows that by increasing the amount of label noise, the accuracy of the SVM classifier model decrease linearly on original dataset; however, by applying our method we obtain high accuracy on the cleansed dataset by identifying noise label instances and then filtering them out from the dataset. It is also important to note that having label noise in the dataset, affects the evaluation part for unforeseen data. However, considering grand truth in simulation shows label noises on unseen data can be detected and currect labels assignd to them with high accuracy.

One potential concern that may arise is that we do not have a ground truth to assess if the model is biased. In the presence of label noise the validation and test accuracy of the model can be decreased by noisy subjects, and likewise it is not possible to estimate generalization metrics without a ground truth [3]. We appeal to the simulation as a proxy for validating our model as it is not possible to know the ground truth in real data. We analyzed our method on the simulated handwriting digit dataset where identification of the label noise is possible since we have the ground truth. We artificially introduced label noise into the simulation dataset by shuffling their labels. Results showed our method successfully identified the correct labels and the performance of the model increased significantly on new cleansed data.

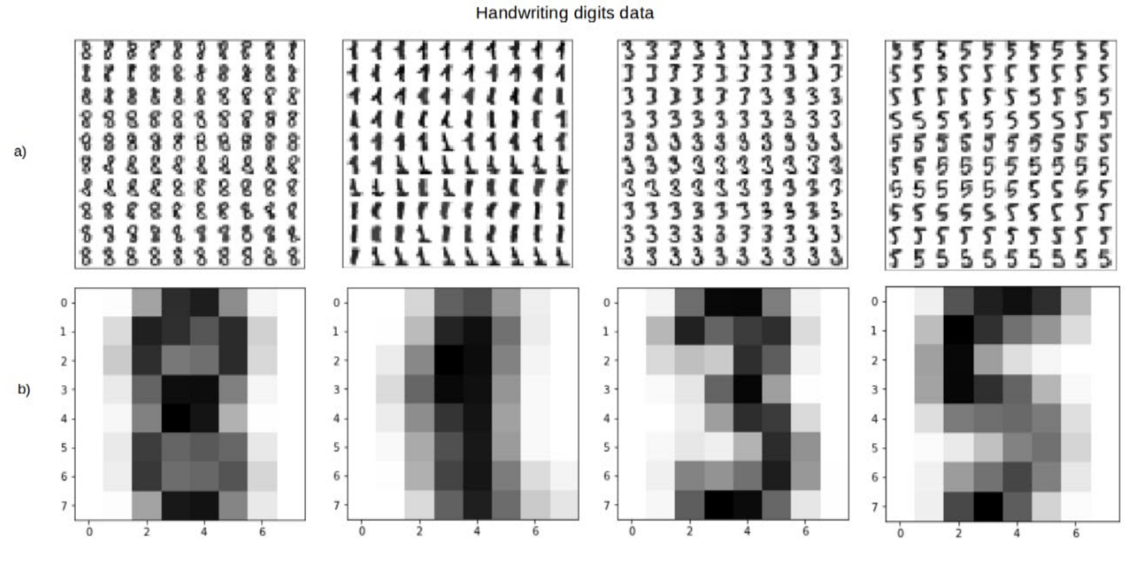


***Figure S1*** *a) Shows random samples from handwriting digits dataset. The numbers of 8,1,3,5 were selected because of their similarities and common features they have; and b) The mean over all samples for each digit.*

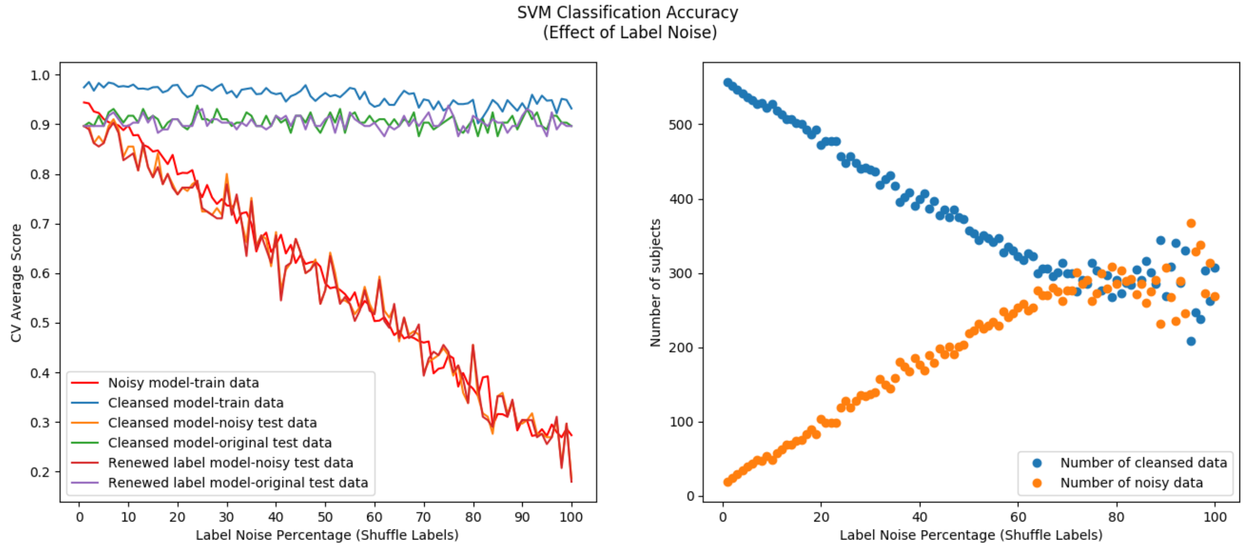


***Figure S2*** *Shows effect of different proportion of label noise on accuracy of SVM classifier and improving accuracy using data cleansing approach. Corrupting data by shuffling labels of instances and adding synthetic label noises to ground truth decreases the performance of the predictive model. By applying data cleansing classification filtering method, we could identify noisy labels and improve the model performance. Even when 80% of labels in dataset are noisy, model accuracy was boosted from 0.38 in the noisy dataset to 0.9 in the cleansed dataset. The effect of label noise on splitting dataset also shows evaluating data cleansins is challenging when the ground truth are not available and may mislead during interpretation.*

**Deep learning:** We analyzed the cleansed dataset with a deep convolutional neural network (CNN) ResNet model. Using a deep learning model, we can circumvent feature engineering which is basically extracting useful representations and compare its result without incorporating an explicit dimension reduction step. The architecture of our model is the modification of the open-source Pytorch implementation of the ResNet framework [4]. 3D structural MRI scans are seen by the convolution layer followed by batch normalization and a rectifier unit layer. The output of these layers goes through the max pooling layer and then goes through the Residual block of containing a series of 3D convolutional, batch normalization and non-linear rectifier (ReLU) layers [5]. The output of residual blocks feeds to into the average pooling layer followed by fully connected layers. The output of this layer is the class probability score. See **Fig S3**.

We analyzed the performance of our deep learning model on the first iteration of classification filtering. The accuracy of our deep supervised convolutional neural network (ResNet) model improved about 20% from 0.65 to 0.79 on an average of 6 binary classification task using DSM-IV labels and by 23% from 0.60 to 0.74 on 6 binary classification task using Biotype labels. **Fig S4** shows the receiver operating characteristic (ROC) plot of these 12 binary classification tasks on our deep model. The boxplot of the accuracy of 12 classification tasks on cleansed and original dataset shows in **Fig S5**.

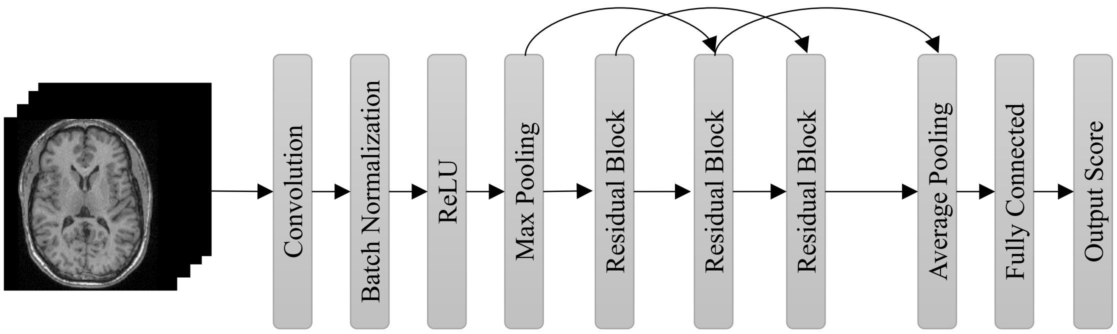


***Fig S3*** *Deep model convolutional neural network ResNet architecture.* 3D structural MRI scans are seen by the convolution layer followed by batch normalization and a rectifier unit layer. The output of these layers goes through max pooling layer and then go through Residual block of containing a series of 3D convolutional, batch normalization and non-linear rectifier (ReLU) layers

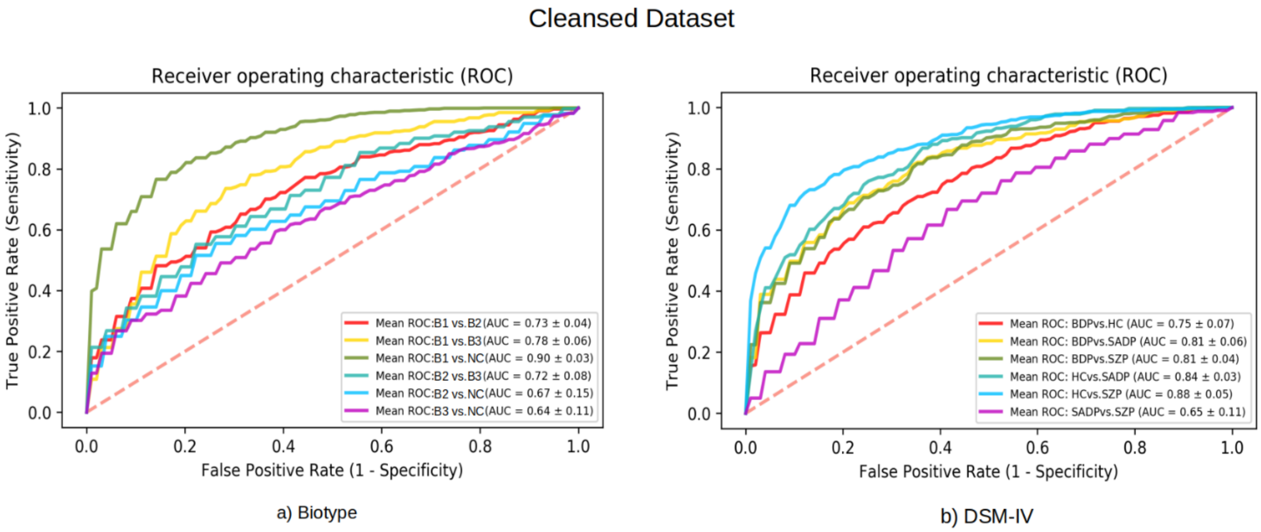


***Figure S4*** *ROC plot of 12 binary classification tasks using Biotype and DSM-IV labels on cleansed dataset after 1st iteration of our data cleansing approach before convergence. After one iteration of classification filtering on data and training deep model using cleansed dataset the accuracy improved about 20% from 0.65 to 0.79 on an average of 6 binary classification task using DSM-IV labels and by 23% from 0.60 to 0.74 on 6 binary classification task using Biotype labels.*

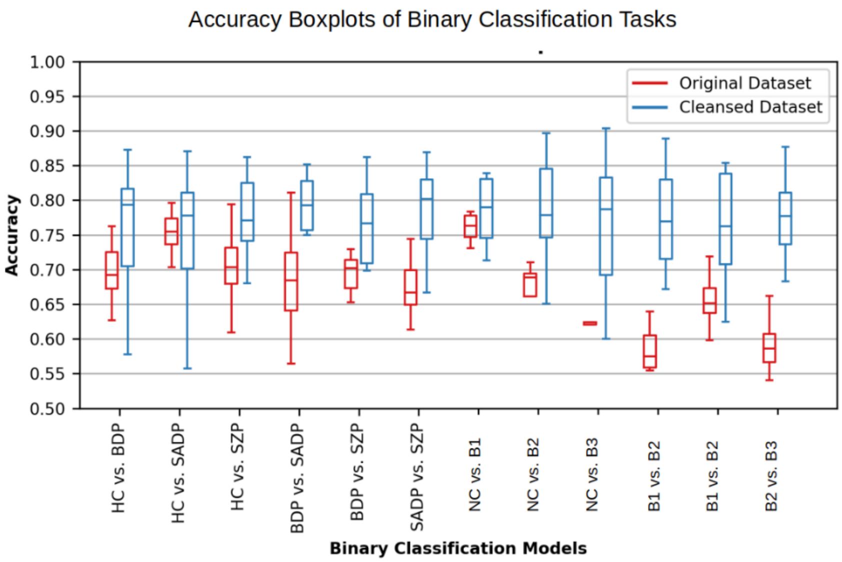


***Figure S5*** *Accuracy boxplot of 12 binary classification tasks on deep model on cleansed dataset after 1st iteration. Red are boxplot accuracy of stratified cross validation using original label. Blue boxplot shows the accuracy of different stratified cross validation run on cleansed dataset after a single classification voting filtering. The figure shows accuracy improved on cleansed dataset after a single iteration of classification filtering.*

### REFRENCES


[1] B. A. Clementz, J. A. Sweeney, J. P. Hamm, E. I. Ivleva, L. E. Ethridge, G. D. Pearlson, M. S. Keshavan and C. A. Tamminga, "Identification of distinct psychosis biotypes using brain-based biomarkers," *American Journal of Psychiatry,* vol. 173, no. 4, pp. 373-384, 2015.

[2] E. I. Ivleva, B. A. Clementz, A. M. Dutcher, S. J. Arnold, H. Jeon-Slaughter, S. Aslan, B. Witte, G. Poudyal, H. Lu, S. A. Meda and others, "Brain structure biomarkers in the psychosis biotypes: findings from the bipolar-schizophrenia network for intermediate phenotypes," *Biological psychiatry,* vol. 82, no. 1, pp. 26-39, 2017.

[3] B. Frenay and M. Verleysen, "Classification in the presence of label noise: a survey," *IEEE transactions on neural networks and learning systems,* vol. 25, no. 5, pp. 845-869, 2013.

[4] A. Paszke, S. Gross, S. Chintala, G. Chanan, E. Yang, Z. DeVito, Z. Lin, A. Desmaison, L. Antiga and A. Lerer, "Automatic differentiation in pytorch," 2017.

[5] A. Abrol, H. Rokham and V. D. Calhoun, "Diagnostic and Prognostic Classification of Brain Disorders Using Residual Learning on Structural MRI Data," in *41st Annual International Conference of the IEEE Engineering in Medicine & Biology Society (EMBC)*, Berlin, 2019.
